# Pupil dilation and the slow wave ERP reflect surprise about choice outcome resulting from intrinsic variability in decision confidence

**DOI:** 10.1101/2020.06.25.164962

**Authors:** Jan Willem de Gee, Camile M.C. Correa, Matthew Weaver, Tobias H. Donner, Simon van Gaal

**Affiliations:** Department of Psychology and Amsterdam Brain & Cognition (ABC), University of Amsterdam, Amsterdam, the Netherlands; Department of Neurophysiology and Pathophysiology, University Medical Center Hamburg-Eppendorf, Hamburg, Germany; Department of Neuroscience, Baylor College of Medicine, Houston, TX, USA; Jan and Dan Duncan Neurological Research Institute, Texas Children’s Hospital, Houston, TX, USA; Centre of Functionally Integrative Neuroscience, Aarhus University, Aarhus, Denmark

**Keywords:** prediction error, confidence, arousal, consciousness, pupil size

## Abstract

Central to human and animal cognition is the ability to learn from feedback in order to optimize future rewards. Such a learning signal might be encoded and broadcasted by the brain’s arousal systems, including the noradrenergic locus coeruleus. Pupil responses and the positive slow wave component of event-related potentials reflect rapid changes in the arousal level of the brain. Here we ask whether and how these variables may reflect surprise: the mismatch between one’s expectation about being correct and the outcome of a decision, when expectations fluctuate due to internal factors (e.g., engagement). We show that during an elementary decision-task in the face of uncertainty both physiological markers of phasic arousal reflect surprise. We further show that pupil responses and slow wave ERP are unrelated to each other, and that prediction error computations depend on feedback awareness. These results further advance our understanding of the role of central arousal systems in decision-making under uncertainty.

## Introduction

Pupil dilation at constant luminance has been related to deviant (unexpected) stimuli (Alamia, VanRullen, Pasqualotto, Mouraux, & Zenon, 2019; Bianco, Ptasczynski, & Omigie, 2020; Kamp & Donchin, 2015a; Kloosterman et al., 2015; Knapen et al., 2016; Liao, Yoneya, Kashino, & Furukawa, 2018; Murphy, Vandekerckhove, & Nieuwenhuis, 2014; Raisig, Welke, Hagendorf, & van der Meer, 2010; Van Slooten, Jahfari, Knapen, & Theeuwes, 2018; Wetzel, Buttelmann, Schieler, & Widmann, 2016; Zhao et al., 2019), behavioral error awareness (Critchley, 2005; Maier, Ernst, & Steinhauser, 2019; Murphy, Boonstra, & Nieuwenhuis, 2016) and to key computational variables such as “uncertainty” about the state the world and/or the right course of action and subsequent “surprise” after finding out (Colizoli et al., 2018; De Berker et al., 2016; Filipowicz et al., 2020; Findling et al., 2019; Krishnamurthy et al., 2017; Lempert et al., 2015; Murphy et al., 2020; Nassar et al., 2012; O’Reilly et al., 2013; Preuschoff et al., 2011; Richer & Beatty, 1987; Urai et al., 2017; Van Slooten et al., 2018; Vincent et al., 2019). Pupil size closely tracks fluctuations in the cortical arousal state mediated by subcortical neuromodulatory systems like the noradrenergic locus coeruleus and the cholinergic basal forebrain (Aston-Jones & Cohen, 2005; Joshi & Gold, 2020; Larsen & Waters, 2018; McGinley et al., 2015). Indeed, during elementary decisions, the role of these neuromodulatory systems might be to broadcast information about momentary uncertainty and surprise (Aston-Jones & Cohen, 2005; Bouret & Sara, 2005; Doya, 2008; Lak, Nomoto, Keramati, Sakagami, & Kepecs, 2017; Parikh, Kozak, Martinez, & Sarter, 2007; Yu & Dayan, 2005), a process that can be read-out by monitoring pupil size.

In this literature, uncertainty and surprise depended lawfully on *external* factors, such as the strength of the stimulus that needs to be discriminated or the volatility of the environment. Strikingly, even when we are given the same information to act on and all external factors are held constant, we will often choose differently each time when asked to make a decision (Glimcher, 2005; Gold & Shadlen, 2007; Sugrue, Corrado, & Newsome, 2005; Wyart & Koechlin, 2016). Such repeated decisions also tend to be associated with varying levels of uncertainty (or the inverse: confidence in being correct) (Fleming & Dolan, 2012; Fleming & Lau, 2014; Meyniel et al., 2015). Choice and confidence variability of this kind must be driven by *internal* variables.

It is unknown if pupil-linked arousal tracks surprise about choice outcome that results from intrinsic variability in decision confidence. This is important because deviations between objective task performance and subjective decision confidence are commonly observed, both in healthy humans as well as in several pathologies. Furthermore, it is currently unclear how peripheral markers relate to neural markers of surprise. The phasic release of neuromodulators may be captured in the size of different components of the late positive complex, the P3a, P3b and slow wave event-related potential (ERP) components, as measured with electroencephalography (EEG) (Boldt & Yeung, 2015; Brown et al., 2015; Friedman et al., 1973; Jepma et al., 2016; Kamp & Donchin, 2015; Murphy et al., 2011; Nieuwenhuis et al., 2005; Pineda et al., 1989; Polich, 2007; Steinhauer & Zubin, 1982). The late positive complex has been shown to scale with novelty, surprise and perceptual confidence in previous studies (Boldt & Yeung, 2015; Yeung & Summerfield, 2012). Finally, it remains an open question if and how surprise also depends on the subjective awareness of the feedback stimulus. Although unconscious stimuli are known affect a plethora of cognitive processes, it is unknown how important feedback awareness is for prediction error computation (van Gaal & Lamme, 2012).

To tackle these questions, we combined an elementary perceptual decision paradigm, including explicit confidence ratings and high or low visibility feedback, with simultaneous pupil size and EEG recordings. We found (i) that both feedback-related pupil responses and ERP slow wave amplitudes reflected surprise about decision outcome, (ii) that the same pupil and ERP amplitudes were unrelated to each other, and (iii) that surprise about decision outcome, as reflected by the pupil and/or slow wave ERP, depends on the conscious access to the feedback stimulus.

## Materials and Methods

### Subjects

Thirty-two students from the University of Amsterdam (23 women; aged 18-24) participated in the study for course credits or financial compensation. All subjects gave their written informed consent prior to participation, were naive to the purpose of the experiments, and had normal or corrected-to-normal vision. All procedures were executed in compliance with relevant laws and institutional guidelines and were approved by the local ethical committee of the University of Amsterdam.

### Tasks

Subjects participated in three experimental sessions, separated by less than one week. We will first explain the main task, performed in session two and three, and thereafter the tasks performed in the first session. In each session, subjects were seated in a silent and dark room (dimmed light), with their head positioned on a chin rest, 60 cm in front of the computer screen. The main task was performed while measuring Pupil and EEG responses.

#### Main task: orientation discrimination task (sessions 2 and 3)

Stimuli were presented on a screen with a spatial resolution of 1280×720 pixels, run at a vertical refresh rate of 100 Hz. Each trial consisted of the following consecutive intervals (**Fig. 1A**): (i) the baseline interval (1.6-2.1 s); (ii) the stimulus interval (0.5 s; interrogation protocol), the start of which was signaled by a tone (0.2 s duration); (iii) the response period (terminated by the participant’s response); (iv) a delay (uniformly distributed between 1.5 and 2 s); (v) the feedback interval (0.5 s), the start of which was signaled by the occurrence of a tone (0.2 s duration); (vi) a delay (uniformly distributed between 1.5 and 2 s); (vii) the feedback identity response period (terminated by the participant’s response).

**Figure 1.**
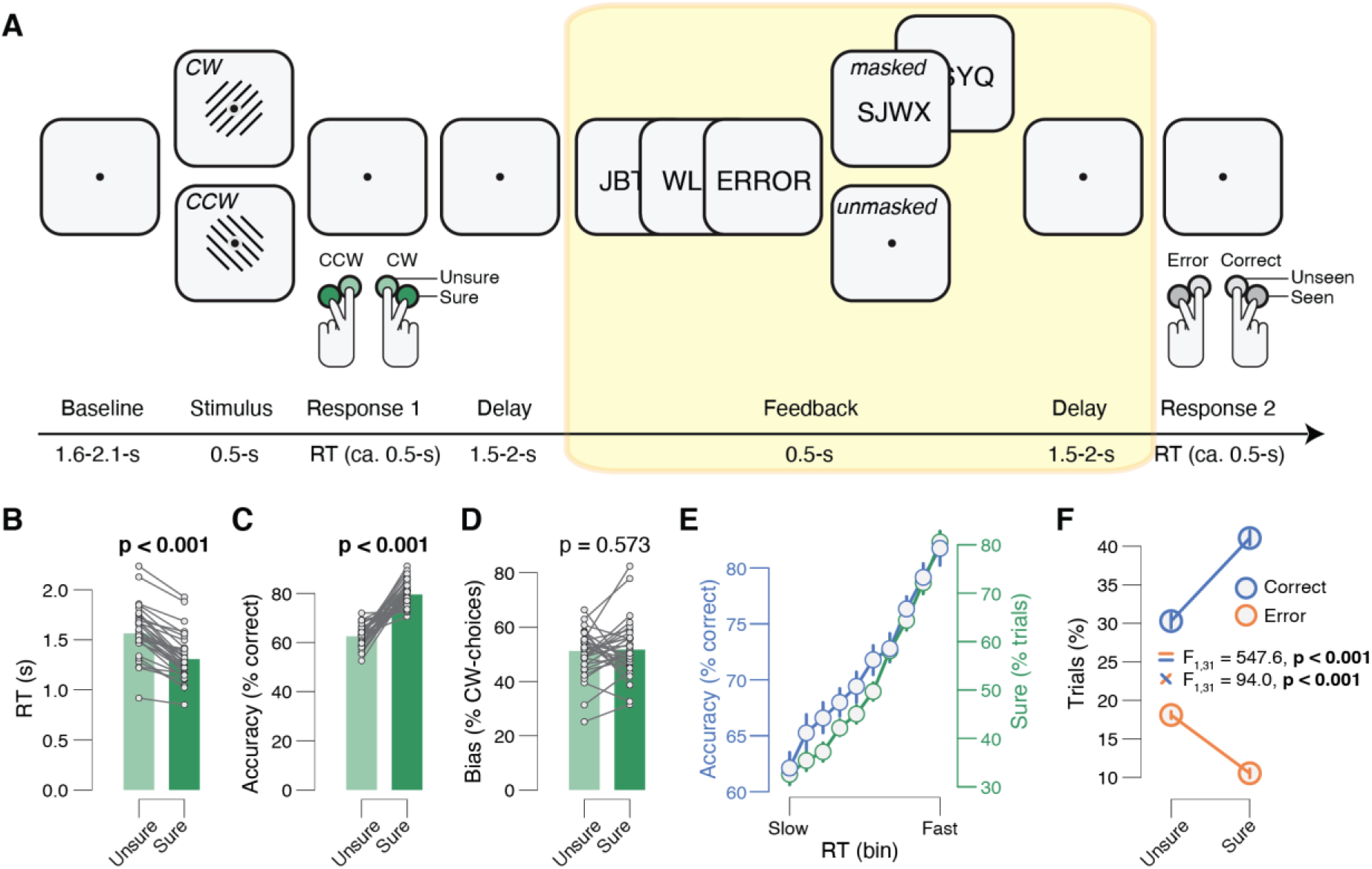
Task and behavior. **(A)** Sequence of events during a single trial. Subjects reported the direction and level of confidence in the decision about a Gabor patch by pressing one of four buttons (CCW=counter-clock-wise, CW=clock-wise; CCW sure; CCW unsure, CW unsure, CW sure). After the decision interval, veridical feedback was presented indicating the correctness of the response. Subjects reported the identity and visibility of the feedback stimulus (the word “error” or “correct” in Dutch) by pressing one of four buttons (seen error; unseen error; unseen correct; seen correct; see Methods for details). **(B)** RT separated for sure vs. unsure decisions. Data point, individual subjects; stats, paired-samples t-test. **(C)** As B, but for accuracy. **(D)** As B, but for choice bias. **(E)** Proportion of correct (blue) and sure (green) choices for 10 RT-defined bins. Error bars, s.e.m. **(F)** Proportion of trials sorted by correctness (error, correct) and confidence (sure, unsure). Stats, ANOVA (Materials and Methods); error bars, s.e.m. Panels B-F: group average (N=32).

During Gabor presentation the luminance across all pixels was kept constant. A sinusoidal grating (1.47 cycles per degree) was presented for the entire stimulus interval. The grating was either tilted 45° (clockwise, CW) or 135° (counter-clockwise, CCW). Grating orientation was randomly selected on each trial, under the constraint that it would occur on 50% of the trials within each block of 60 trials. The grating was presented in a Gaussian annulus of 11.4 cm, with a 10.85 degrees visual angle (1.47 cycles per degree). Feedback was signaled by the Dutch word “goed” (correct feedback) or the word “fout” (incorrect feedback), from now on referred to as “correct” and “error” feedback. The words were presented for three frames just below fixation. Feedback was either masked, by presenting both forward as well as backward masks (masks1-masks2-feedback-masks3-masks4) or unmasked, by presenting only forward masks (masks1-masks2-feedback). Each mask consisted of 6 randomly scrambled letters (without the letters making up the words “goed” or “fout”). Masks’ types were presented two frames each. Feedback type (masked vs. unmasked) was randomly selected on each trial, under the constraint that it would occur on 50% of the trials within each block of 60 trials (**Fig. 1A**).

Throughout the main experiment, the contrast of the Gabor was fixed at the individual threshold level that yielded about 70% correct choices. Each subject’s individual threshold contrast was determined before the main experiment using an adaptive staircase procedure (Quest). The corresponding threshold contrasts yielded a mean accuracy of 70.9% correct (±0.44 % s.e.m.) in the main experiment.

Subjects performed between 12 and 17 blocks (distributed over two measurement sessions), yielding a total of 720–1020 trials per participant. Subjects were instructed to report the orientation of the Gabor, and simultaneously their decision confidence in this decision, by pressing one of four response buttons with their left or right index or middle finger: left middle finger: CCW, sure; left index finger: CCW, unsure; right index finger: CW, unsure; right middle finger: CW, sure. Subjects were also instructed to report the identity and visibility of the feedback by pressing one of four response buttons with their left or right index or middle finger: left middle finger – “error”, seen; left index finger – “error”, unseen; right index finger – “correct”, unseen; right middle finger – “correct”, seen. For analyses, we defined high visibility feedback trials as trials on which the feedback was unmasked and subjects reported it as “seen”. We defined low visibility feedback trials as trials on which feedback was masked and subjects reported it as “unseen”.

Note that indeed, the masking procedure revealed two clear categories: on unmasked trials subjects were 99.82% correct in their discrimination between feedback identity (error/correct) (s.e.m.=0.05%) whereas in the masked condition they were 71.21% correct (s.e.m.=1.69%). Note that chance level in the feedback discrimination task is not 50%, because overall Gabor discrimination performance was 70.9% and feedback presentation was veridical. Therefore, subjects may have been able to anticipate the likelihood of being correct/wrong.

#### Passive viewing task (session 1)

In this control task, subjects fixated their gaze at the center of the screen and passively viewed the words “goed” (correct) and “fout” (error), randomly presented for 100 times. Words were presented for three frames (100 Hz refresh rate) and were not masked.

#### Forced-choice visibility task (session 1)

In this control task, the words “goed” (correct) or “fout” (error) were presented in the same way as in the main experiment (see above), that is, in a masked or unmasked manner (same timings and presence or absence of masks as described above). Subjects were instructed to report the identity of the presented words, by pressing one of two response buttons with their left or right index finger: left – “error”; right – “correct” (the stimulus-response mapping was counter-balanced across trials, and was indicated on the screen after each trial). Subjects performed two blocks, yielding a total of 200 trials per participant.

In total we tested 49 subjects in the first behavioral and eye-tracking session (namely, the passive viewing and forced-choice visibility tasks), considered a pre-screening procedure. We invited 32 subjects to main experiment performed in the second and third session. Six subjects did not enter the main experiment due to various reasons (e.g. drop-out, extensive blinking (subjectively assessed by the experimenter during the first session, not based on formal analyses)). Of the remaining 43 subjects, the 32 subjects with the lowest discrimination performance score on the forced-choice discrimination task were invited for the second and third session (resulting in an accuracy cut-off of > 73%). Discrimination performance for the 32 included subjects varied between 49% and 73% correct. Included subjects were on average 98.87% (SEM=0.02) correct in the unmasked condition and 61.9% (SEM=0.02) correct in the masked condition. The average percentage of correct responses for masked words exceeded chance-level performance (t_31_=11.26, p<0.001).

#### Priming task (session 1)

In this control task, subjects were instructed to respond as fast and accurately as possible to eight Dutch words, randomly selected out of five of positive (laugh, happiness, peace, love, fun) and 5 (death, murder, angry, hate, war) of negative in valence, by pressing one of two response buttons with their left or right index finger: left – negative; right – positive. Unknown to our subjects, these words were preceded by the masked words “goed” and “fout”, respectively “correct” and “incorrect”, three frames each before the positive or negative word targets (12 frames each) in 100 Hz refresh rate. This yielded congruent and incongruent trials. Subjects performed six blocks, yielding a total of 480 trials per participant.

### Data acquisition

The diameter of the left eye’s pupil was tracked at 1000 Hz with an average spatial resolution of 15–30 min arc, using an EyeLink 1000 system (SR Research, Osgoode, Ontario, Canada). EEG data was recorded and sampled at 512 Hz using a BioSemi Active Two system. Sixty-four scalp electrodes were distributed across the scalp according to the 10–20 International system and applied using an elastic electrode cap (Electro-cap International Inc.) Additional electrodes were two electrodes to control for eye-movements (left eye, aligned with the pupil, vertically positioned, each referenced to their counterpart), two reference electrodes at the ear lobes to be used as reference and two electrodes for heartbeat (positioned at the left of the sternum and in the right last intercostal space).

### Data analysis

#### Eye data preprocessing

Periods of blinks and saccades were detected using the manufacturer’s standard algorithms with default settings. The subsequent data analyses were performed using custom-made Python software. The following steps were applied to each pupil recording: (i) linear interpolation of values measured just before and after each identified blink (interpolation time window, from 150 ms before until 150 ms after blink), (ii) temporal filtering (third-order Butterworth, low-pass: 10 Hz), (iii) removal of pupil responses to blinks and to saccades, by first estimating these responses by means of deconvolution, and then removing them from the pupil time series by means of multiple linear regression (Knapen et al., 2016), and (iv) conversion to units of modulation (percent signal change) around the mean of the pupil time series from each block.

#### Quantification of feedback-evoked pupillary responses

We computed feedback-evoked pupillary response amplitude measures for each trial as the mean of the pupil size in the window 0.5 s to 1.5 s from feedback, minus the mean pupil size during the 0.5 s before the feedback. This time window was chosen to be centered around the peak of the pupil response to a transient event (like the feedback in our task; **Fig. 2A**) (de Gee, Knapen, & Donner, 2014; Hoeks & Levelt, 1993).

**Figure 2.**
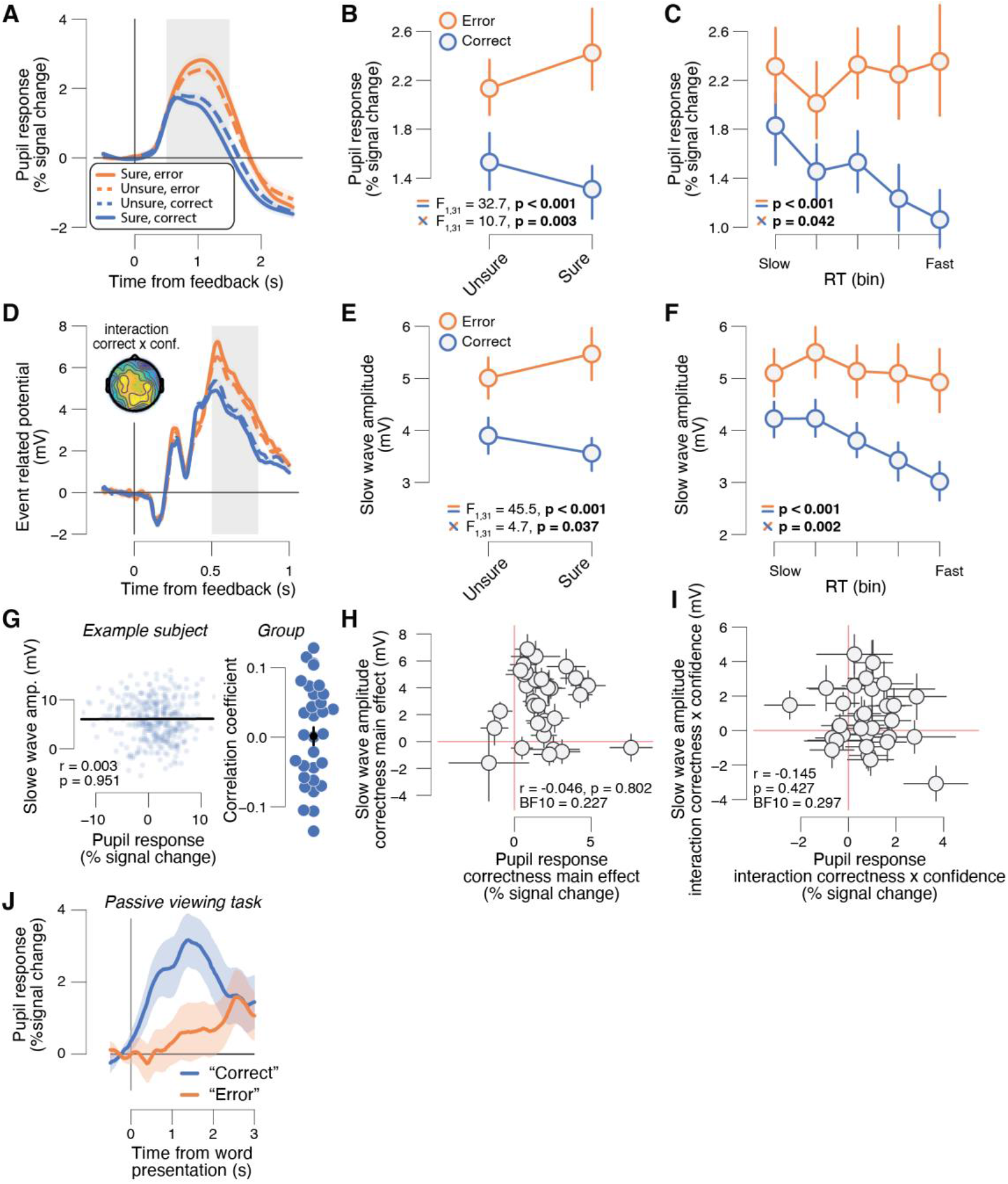
Feedback-related pupil response and slow wave amplitudes report a prediction error. **(A)** High visibility feedback-related pupil time course, sorted by correctness (error, correct) and confidence (sure, unsure). Grey box, interval for averaging pupil response values on single trials. **(B)** High visibility feedback-related pupil responses sorted by correctness (error, correct) and confidence (sure, unsure). Stats, ANOVA (Materials and Methods): top, correctness main effect; bottom, correctness x confidence interaction. **(C)** High visibility feedback-related pupil responses sorted by correctness (error, correct) and RT. Stats, mixed linear model (Materials and Methods). **(D)**, As A, but for the high visibility feedback event-related potential (ERP) time courses. Head map, correctness x confidence interaction (map limits [−1 1]). **(E,F)**, as B,C but for the high visibility feedback-related slow wave amplitudes. **(G)** Left: trial-by-trial relationship between feedback-related pupil responses and slow wave amplitudes for one example subject. Data points represent single trials. Right: correlation coefficient of the same relationship separately for each subject. Data points, single subjects (green dots represent significant correlations [p<0.05]); black dot with error bars, group average ± s.e.m. **(H)** High visibility feedback-related slow wave amplitude correctness main effect ([error/unsure + error/sure] - [correct/unsure + correct/sure]) plotted against feedback-related pupil response correctness main effect. Stats, Pearson correlation; datapoints, individual subjects (N=32); error bars, 60% confidence intervals (bootstrap). **(I)** As H, but for correctness x confidence interaction effects ([error/sure - correct/sure] - [error/unsure - correct/unsure]). **(J)** High visibility event related pupil time course sorted by the words “correct” and “error” during a passive viewing experiment (Materials and Methods). All panels except H,I: group average (N=32); shading or error bars, s.e.m.

It is commonly observed that task-evoked pupil responses are negatively correlated to pre-trial baseline pupil size (de Gee et al., 2014; Filipowicz, Glaze, Kable, & Gold, 2020; Gilzenrat, Nieuwenhuis, Jepma, & Cohen, 2010; Mridha et al., 2019), which is partly due to floor and ceiling effects and a general reversion to the mean. Indeed, pupil responses that occurred time-locked to the decision about Gabor orientation depended negatively on pre-trial baseline pupil size: the group average correlation coefficient (± s.e.m.) was −0.314 (0.015). However, the feedback-related pupil responses (that occurred later in the trial) were not correlated to pre-trial baseline pupil size: the group average correlation coefficient (± s.e.m.) was 0.037 (0.079) in the high visibility condition), and 0.046 (0.061) in the low visibility condition (see **Figs. S1** **&** **S2** for individual subject scatter plots). Therefore, we did not include pre-trial baseline pupil size as a covariate in any of our analyses. All results remain qualitatively the same after including pre-trial baseline pupil size as a covariate (data not shown).

#### EEG data preprocessing

Standard pre-processing steps were performed in EEGLAB toolbox in Matlab. Data were bandpass filtered from 0.1 to 40 Hz off-line for ERP analyses. Epochs ranging from −1 to 2 seconds around feedback presentation were extracted. Linear baseline correction was applied to these epochs using a −200 to 0 ms window. The resulting trials were visually inspected and those containing artifacts (e.g. movement artifacts) were removed manually. Moreover, electrodes that consistently contained artifacts were interpolated, entirely or per bad epoch. Finally, using independent component analysis (ICA), artifacts caused by blinks, horizontal eye-movements, heartbeats and single electrode noise (when excessive only in short periods of time, otherwise the entire electrode was removed and interpolated) were automatically removed from the EEG data using EEGlab. We took a conservative approach and only removed ICA components that were clearly not related to brain activity. On average 3.95 components were removed per subject.

#### Quantification of slow wave component of the feedback-related ERP

We focused on the slow wave component of the feedback-related ERP. The slow wave amplitude on each trial was defined as the mean electrophysiological response in the window 0.5 to 0.8 seconds after feedback presentation, measured in a central region of interest (ROI): the averaged signal of electrodes F1, Fz, F2, FCz, FC1, FC2, Cz, C1, C2, CPz, CP1, CP2, Pz, P1, P2. The region of interest (electrodes) as well as time-window of interest for the single-trial slow waves (0.5 to 0.8 sec) were a priori selected and were identical to a previous study of our group on a similar topic (Correa et al., 2018). ERPs were calculated by taking the mean across all trials. Note that the selection of a region of interest in space (electrodes) in combination with a specific time-window for the EEG data ensures that the analysis protocol becomes highly similar between pupil and ERP responses. In fact, the same ANOVA’s can be performed with exactly the same factors for both measures (correctness, confidence, visibility), which makes the pupil and ERP results directly comparable and more intuitive to interpret.

For exploratory analysis on the feedback-related negativity (FRN) we used exactly the same region of interest as for the slow wave, again identical to a previous study (Correa et al. 2018). The FRN peaked around 400 ms at central electrodes, similar to Correa et al (2018). We used 350-450 ms post outcome stimulus as our time window of interest for the ANOVA’s.

#### Behavioral analyses

Behavioral and statistical analyses were performed in Python. We excluded trials in which a subject blinked during the presentation of the Gabor stimulus (duration, 0.5 s) or the timepoints used to compute feedback-related pupil responses and slow wave amplitudes (0.5 – 1.5 s from feedback). The group average (± S.E.M) trial-wise blink rate was 15.43% (2.85%). We only considered unmasked trials reported as seen as high visibility feedback (99% of unmasked trials) and masked trials indicated as unseen as low visibility feedback (89.2% of masked trials). The results are qualitatively the same when using all trials and splitting on seen vs. unseen (irrespective of masking), or when using all trials and splitting on masked vs. unmasked (irrespective of subjective visibility report) (data not shown). RT was defined as the time from stimulus offset until the button press.

Autocorrelations in performance might give rise to an artificial correlation between feedback-related pupil responses or slow wave amplitude and behavioral performance on the subsequent trial. Indeed, during the course of an experiment autocorrelation is typically observed in RTs and accuracy (Dutilh et al., 2012; Gilden, 2001; Palva et al., 2013). This could be due to slow drifts in behavioral state factors (e.g. motivation, arousal, attention). We reasoned that a prediction error signal cannot affect performance on the previous trial (because of temporal sequence); instead, any observed relationship must be due to slow autocorrelations in performance. Therefore, in order to isolate the impact of rapid (trial-by-trial) prediction error signal on performance on the next trial from slow ongoing fluctuations, we took the *difference* between the correlation coefficients that captured the trial-by-trial relationship between a prediction error signal (pupil or slow wave amplitude) and performance on the next trial versus the previous. A similar approach has been used before (Desender, Boldt, Verguts, & Donner, 2019).

#### Statistical comparisons

We used 2×2 repeated measures ANOVA to test for the main effect of being correct, and for the interaction effect between correctness and confidence. With a 2×2×2 repeated measures ANOVA we tested whether these main and interaction effects were different between the high and low visibility conditions. We used mixed regression modeling to quantify the trial-by-trial statistical dependence of feedback-related pupil responses or slow wave amplitudes on RT and accuracy. Error variance caused by between-subject differences was accounted for by adding random slopes to the model. Random slopes for a given factor (RT or accuracy) were added only when this increased the model fit, as assessed by model comparison using Bayesian Information Criterion (BIC). We used Pearson correlation to quantify the within- and across-subject correlations. We used the paired-samples t-test to test for differences in RT, accuracy or choices between sure and unsure trials, and between congruent and incongruent priming conditions.

#### Data and code sharing

The data are publicly available on [to be filled in upon publication]. Analysis scripts are publicly available on [to be filled in upon publication].

## Results

### A prediction error signature in behavior

During simultaneous pupillometry and EEG recordings, thirty-two human subjects performed a challenging contrast orientation discrimination task (three experimental sessions per subject, on different days). On each trial this involved discriminating the orientation (clockwise [CW] vs. counter-clockwise [CCW]) of a low-contrast Gabor, explicit confidence ratings and feedback (**Fig. 1A**). The Gabor’s contrast was adjusted individually such that each subject performed at about 70% correct (**Fig. 1C**; Materials and Methods). Subjects simultaneously indicated their CW vs. CCW-choice and the accompanying confidence in that decision (sure vs. unsure; type 1 confidence (Galvin, Podd, Drga, & Whitmore, 2003), see **Fig. 1A**). These explicit ratings provided a window into the trial-to-trial fluctuations of decision confidence, which may shape prediction error signals after decision outcome (feedback), and physiological correlates thereof. The Dutch words for “error” or “correct” provided feedback about the correctness of the preceding CW vs. CCW-choice. The feedback was masked by random letters on 50% of trials. This was done to manipulate feedback awareness, which may in turn affect uncertainty (about feedback valence) and phasic measures of central arousal state. At the end of the trial, subjects had to indicate the subjective visibility and identity (the word “error” or “correct”) of the feedback stimulus (**Fig. 1A**). This allowed us to post-hoc sort trials based on the combination of masking strength and subjective visibility (Materials and Methods).

Subjects’ choice behavior indicated that they successfully introspected perceptual performance: subjects were faster and more accurate when they were confident in their decision (**Fig. 1B,C**), a typical signature of confidence (Kamp & Donchin, 2015b; Meyniel, Sergent, Rigoux, Daunizeau, & Pessiglione, 2013). There was no relationship between confidence and decision bias (**Fig. 1D**). In line with earlier work (Sanders, Hangya, & Kepecs, 2016; Urai et al., 2017), reaction times (RTs) predicted accuracy and confidence, with more accurate and confident choices for faster RTs (**Fig. 1E**). Taken together, these results suggest that subjects in our task were able to introspect perceptual performance well.

Negative feedback (‘error’) was more surprising than positive feedback (‘correct’), because subjects performed well above chance (~71% correct). Negative feedback should be especially surprising when subjects were relatively sure about the correctness of the preceding choice. In contrast, positive feedback should be least surprising when they were relatively sure about the correctness of the preceding choice. In line with this intuition, trial counts followed the expected ordering (from least to most often, that is, from most to least surprising): sure/error, unsure/error, unsure/correct, sure/correct (**Fig. 1F**). For trial counts, there was thus a significant main effect for correctness ([error/unsure + error/sure] - [correct/unsure + correct/sure]) and an interaction effect between correctness and confidence ([error/sure - correct/sure] - [error/unsure - correct/unsure]); **Fig. 1F**). Any physiological variable that encodes a prediction error (surprise about decision outcome) should follow a similar pattern. The main effect of correctness may partly reflect error monitoring (Cohen, Wilmes, & van de Vijver, 2011; Ullsperger, Fischer, Nigbur, & Endrass, 2014). Since error monitoring can be triggered purely by the type of feedback received (error vs. correct), we consider especially the interaction between confidence and correctness a signature of a prediction error.

### A feedback-related pupil response and the slow wave component of the feedback related ERP report a prediction error

We tested whether feedback-related pupil responses reflect a prediction error after clearly visible feedback. Using the same 2×2 ANOVA logic, we found that indeed feedback-related pupil responses (to high visibility feedback) were larger for error versus correct feedback (main effect of correctness) and that this error vs. correct difference was larger when subjects’ were sure vs. unsure (interaction correctness x confidence; **Fig. 2A,B**). The slow wave component of the feedback-related ERP exhibited a similar functional profile: slow wave amplitudes were larger after error vs. correct feedback and this effect interacted with decision confidence (**Fig. 2D,E**; see head map in **Fig. 2D** for a topographical distribution of the interaction between confidence and correctness).

We used the reaction times, a sensitive implicit measure of confidence (**Fig. 1E**), to visualize and quantify the pupil- and slow wave-reported prediction errors in a more fine-grained fashion (Braun, Urai, & Donner, 2018; Urai et al., 2017). With a mixed linear model, we quantified the trial-by-trial dependence of the feedback-related pupil response on type of feedback (correct vs. error), RT and their interaction (Materials and Methods). The feedback-related pupil responses were larger for error compared to correct feedback and this effect interacted with RT (**Fig. 2C**). Likewise, the feedback-related slow wave amplitudes were larger for error compared to correct feedback and this effect interacted with RT (**Fig. 2F**).

One influential account (Nieuwenhuis et al., 2005) postulates that the pupil response and the slow wave component of the ERP (P3) are driven by the same central (e.g., neuromodulatory) process. This would predict that both measures not only exhibit the same functional profile on average (**Fig. 2A-F**), but that they are also correlated at the single trial level within subjects, and that the magnitude of their respective main and interaction effects are correlated across subjects. Our data did not provide any evidence for those associations: feedback-related pupil responses and slow wave amplitudes were not correlated at the single trial level within subjects (**Fig. 2G**, group average r=0.001, s.e.m=0.013), and the magnitude of their respective main and interaction effects were not correlated across subjects (**Fig. 2H,I**). This suggests that pupil responses and the slow wave component of ERPs are driven by distinct neural processes, both of which are sensitive to decision confidence and prediction errors.

The subject-wise correctness main effect and correctness x confidence interaction effect of feedback-related pupil responses and slow wave amplitudes were not correlated to subjects’ mean accuracy, mean confidence, or metacognitive sensitivity (meta-d’ [(Maniscalco & Lau, 2012; Verbruggen & Logan, 2009)]) (**Fig. S3**).

We verified that the correctness main effect was not driven by any low-level stimulus characteristics, such as luminance, or the intrinsic valence of the words used as feedback (e.g. being of positive/negative valence; Materials and Methods). To that end, before the main experiment, subjects passively viewed the same feedback stimuli while we measured their pupil size (Materials and Methods). In this passive context, the pupil dilated more after the word “correct” compared to “error” (**Fig. 2J**), which is the opposite of what we found in the main experiment (**Fig. 2A**).

The ERP analyses reported so far were performed on the slow wave ERP component (500-800 ms after feedback). We additionally explored the feedback-related negativity (FRN), a frontocentral negative ERP component associated with choice outcome processing and prediction error computation (Cohen et al., 2011; Correa et al., 2018). The FRN can be observed as a small negative difference for the contrast error minus correct feedback, peaking around 400 ms (**Fig. 2D**; see Correa et al 2018 for a similar timing). Indeed, the FRN was robust after high visibility feedback; however, this effect did not interact with confidence (**Fig. S4A,B**). Finally, we also zoomed in on the peak of the P3 ERP component (500-600 ms after feedback), and the effects were similar to those for the slow wave component (**Fig. S4A,C**).

Taken together, we conclude that both physiological variables, feedback-related pupil responses and the slow wave component of feedback-related ERPs (including the P3), report a prediction error, when feedback is presented fully consciously.

### Physiological correlates of prediction errors depend on feedback awareness

We tested whether feedback-related pupil responses and the slow wave component of feedback-related ERPs also report a prediction error after low visibility feedback (Materials and Methods). We did not find evidence for this. For the feedback-related pupil responses, there was a significant correctness main effect, but no correctness x confidence interaction effect (**Fig. 3A,B**). For the feedback-related slow wave amplitudes, there was no significant main effect of correctness nor an interaction effect thereof with confidence **Fig. 3D,E**). The same was true for the FRN and for the peak of the P3 ERP component (**Fig. S5**).

**Figure 3.**
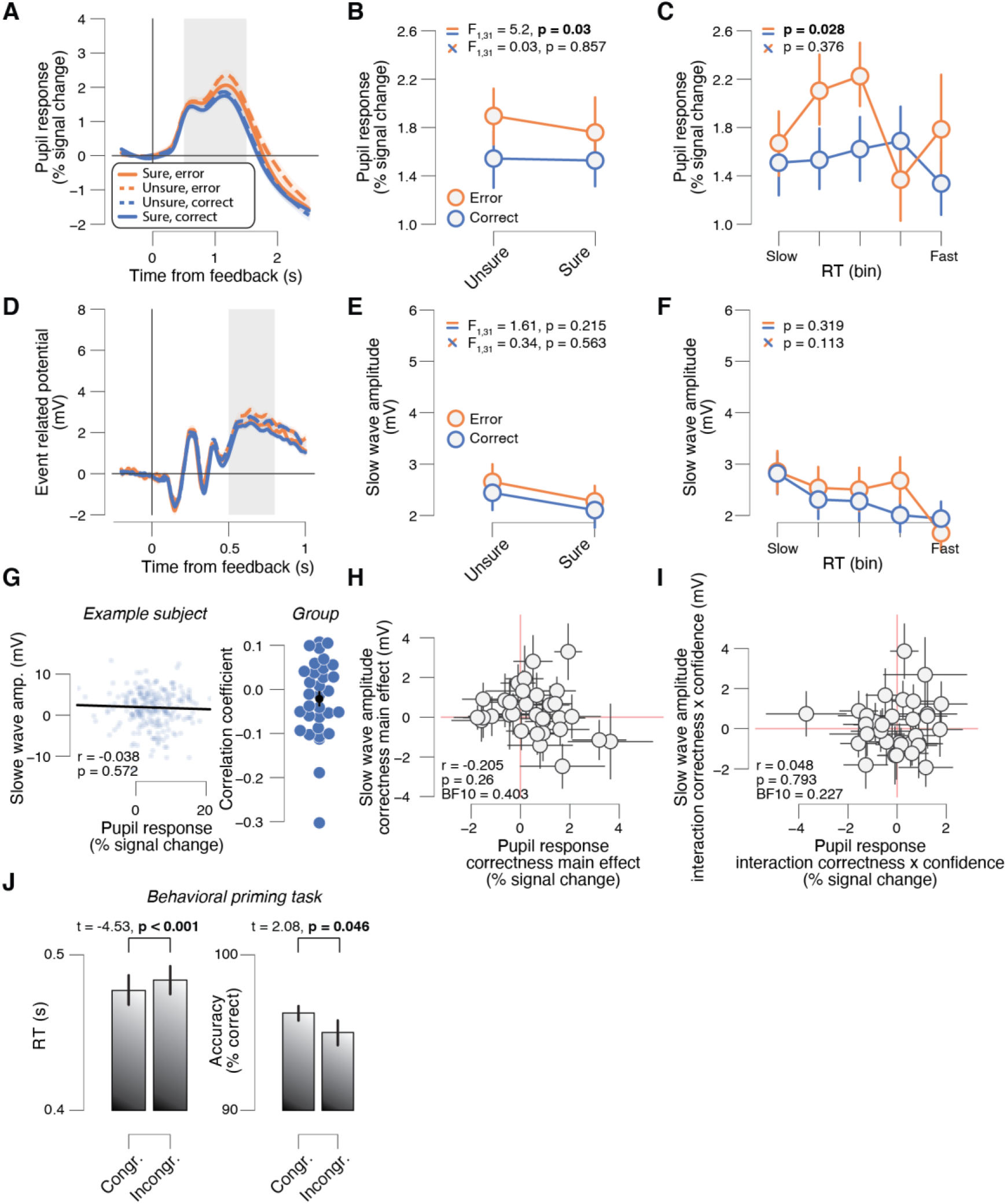
Physiological correlates of prediction errors depend on feedback awareness. **(A)** Low visibility feedback-related pupil time course, sorted by correctness (error, correct) and confidence (sure, unsure). Grey box, interval for averaging pupil response values on single trials. **(B)** High visibility feedback-related pupil responses sorted by correctness (error, correct) and confidence (sure, unsure). Stats, ANOVA (Materials and Methods): top, correctness main effect; bottom, correctness x confidence interaction. **(C)** Low visibility feedback-related pupil responses sorted by correctness (error, correct) and RT. Stats, mixed linear model (Materials and Methods). **(D)**, As A, but for the high visibility feedback event-related potential (ERP) time courses. Head map, correctness x confidence interaction (map limits [−1 1]). **(E,F)**, as B,C but for the low visibility feedback-related slow wave amplitudes. **(G)** Left: trial-by-trial relationship between feedback-related pupil responses and slow wave amplitudes for one example subject. Data points represent single trials. Right: correlation coefficient of the same relationship separately for each subject. Data points, single subjects (green dots represent significant correlations [p<0.05]); black dot with error bars, group average ± s.e.m. **(H)** Low visibility feedback-related slow wave amplitude correctness main effect ([error/unsure + error/sure] - [correct/unsure + correct/sure]) plotted against feedback-related pupil response correctness main effect. Stats, Pearson correlation; datapoints, individual subjects (N=32); error bars, 60% confidence intervals (bootstrap). **(I)** As H, but for correctness x confidence interaction effects ([error/sure - correct/sure] - [error/unsure - correct/unsure]). **(J)** Reaction times (left) and accuracy (right) sorted by congruency (congruent, incongruent) showing typical behavioral priming effects (Materials and Methods). All panels except H,I: group average (N=32); error bars, s.e.m.

We did also not find evidence for a prediction error using the more fine-grained framework based on RTs. Again, for the feedback-related pupil responses, there was a significant effect of feedback type (correct vs. error), but no interaction effect thereof with RT (**Fig. 3C**). For the feedback-related slow wave amplitudes, there was no significant effect of feedback type nor an interaction effect thereof with RT (**Fig. 3F**).

As before, both physiological measures were not correlated at the single trial level within subjects (**Fig. 3G**, group average r=−0.021, s.e.m=0.016), and the magnitude of their respective main and interaction effects were not correlated across subjects (**Fig. 3H,I**).

We verified that the low visibility feedback was not too weak (because of masking) to drive a potential prediction error. To that end, before the main experiment, subjects completed a behavioral priming experiment with the same stimuli and stimulus timings (Materials and Methods). Subject showed typical priming effects: they were faster and more accurate for congruent prime-target pairs versus incongruent pairs (**Fig. 3J**). Thus, the low visibility feedback was not too weak to potentially drive a prediction error.

Feedback-related pupil responses and slow wave amplitudes reported prediction errors significantly better after high versus low visibility feedback. For the feedback-related pupil responses, there was a significant visibility x correctness interaction effect (F_1,31_=11.02, p=0.002) and a visibility x correctness x confidence interaction effect (F_1,31_=6.13, p=0.020). Likewise, for the feedback-related slow wave amplitudes, there was a significant visibility x correctness interaction effect (F_1,31_=34.73, p<0.001). However, there was no significant visibility x correctness x confidence interaction effect (F_1,31_=2.67, p=0.112).

In sum, feedback-related pupil responses and the slow wave component of feedback-related ERPs only reflected a prediction error after high visibility feedback (interaction correctness x confidence). Our results indicate that full visibility of decision feedback is critical to drive a prediction error response.

### High visibility feedback-related slow wave amplitude, but not pupil response, predicts speed-accuracy adaptation on the next trial

Finally, we explored the impact of the observed prediction error responses (feedback-related pupil or slow wave amplitude) on trial-by-trial adjustments of decision-making. To this end, we quantified the relationship between prediction error responses and performance on the subsequent trial, over and above slow drifts in of those variables across trials (see Materials and Methods for details). A correlation between prediction error responses on one trial and performance on the next trial might reflect slow drifts in each of those variables (across multiple trials) as well as the effect of interest here: a rapid (trial-by-trial) adjustment of subsequent decision-making controlled by the prediction error responses. In contrast, such a correlation between a prediction error response on one trial and performance on the *previous* trial cannot reflect adjustments governed by prediction error responses (because of temporal sequence). Therefore, the *difference* between the above two correlations should isolate the effect of prediction error responses on subsequent performance (see also Desender et al., 2019 for a similar approach) and we used this difference measure as our readout of the functional impact of prediction error responses on subsequent performance (accuracy and reaction time).

High visibility feedback-related slow wave amplitudes predicted both, slower and more accurate responses on the subsequent trial (bars in **Fig. S6C,D**), a hallmark of behavioral speed-accuracy adaptation (Cohen & van Gaal, 2013; Desender et al., 2019; Dutilh et al., 2012; Heitz, 2014). These effects were not significant for low visibility feedback (data not shown). By contrast, we found no effect of high visibility feedback-related pupil responses on subsequent performance (bars in **Fig. S6A,B**). High visibility feedback-related slow wave amplitudes did not predict subsequent effects on confidence (group average Δr=−0.126, p=0.161), choice bias (group average Δr=0.145, p=0.278), or choice repetition probability (group average Δr=−0.217, p=0.089). The same was true for low visibility feedback-related slow wave amplitudes (group average Δr=0.023, p=0.808; group average Δr=0.042, p=0.911; group average Δr=−0.116, p=0.443), high visibility feedback-related pupil responses (group average Δr=0.004, p=1.0; group average Δr=−0.01, p=0.97; group average Δr=0.068, p=0.513) and low visibility feedback-related pupil responses (group average Δr=0.085, p=0.369; group average Δr=0.141, p=0.34; group average Δr=−0.079, p=0.369). In sum, high visibility feedback-related slow wave amplitude, but not pupil response magnitude, predicts speed-accuracy adaptation on the next trial.

## Discussion

Pupil responses and the positive slow wave component of ERPs reflect rapid changes in the arousal level of the brain. We investigated whether and how these variables reflect surprise: the mismatch between one’s expectation about being correct and the outcome of a decision, when expectations fluctuate due to internal factors (e.g., engagement). We show that in an elementary decision-task, feedback-related pupil responses and the slow wave component of feedback-related ERPs reflect surprise. We further show that, within and across subjects, pupil responses and the slow wave component of ERPs are unrelated to each other, and that prediction error computations depend on feedback awareness. The results could not be explained by any low-level stimulus characteristics, such as luminance, or the intrinsic valence of the words used as feedback (e.g. being of positive/negative valence, **Fig. 2I**). The reported findings advance our current knowledge about how arousal-linked prediction error computations interact with decision confidence and conscious awareness in in several important ways.

For the first time (to our knowledge), we reveal that pupil responses and the slow wave component of ERPs reflect a prediction error that results from intrinsic variability in subjective decision confidence (with all external variables held constant, e.g., task difficulty). Previous studies have revealed that pupil dilation reflects decision uncertainty and prediction error computation during perceptual choices (Colizoli et al., 2018; Joshi & Gold, 2020; Urai et al., 2017) when task difficulty was manipulated. In these studies, humans performed a random dot motion task incorporating easy and difficult trials depending on the strength of motion coherence. Pupil responses were larger for performance feedback informing that the decision was erroneous vs. correct and this effect was modulated by trial difficulty, in such a way that the pupil dilated most for erroneous decisions based on strong evidence (strong prediction error) and least for correct decisions based on strong evidence (no prediction error). However, several studies have shown that subjective reports of decision confidence do not necessarily track experimental manipulations of task difficulty, for example because confidence estimations are biased due to individual differences in sensitivity to evidence strength or affective value (Fleming & Lau, 2014; Lebreton et al., 2018; Zylberberg et al., 2014). Because we interrogated subjective decision confidence on every single trial, this allowed us to perform post-hoc trial sorting based on trial-by-trial fluctuations in confidence under equal task settings. Thereby we were able to link feedback processing directly to subjective confidence estimations, establishing direct evidence for prediction error computation reported by the pupil and the slow wave component of the ERP.

Our finding that feedback-related pupil responses and the slow wave component of ERPs were uncorrelated within and across subjects is in line with previous studies (Hong, Walz, & Sajda, 2014; Kamp & Donchin, 2015b; Mückschel, Chmielewski, Ziemssen, & Beste, 2017; Murphy et al., 2011; LoTemplio, Silcox, Federmeier, & Payne, 2019) and suggests that these established markers of phasic arousal are driven by distinct sources. Recent animal studies have revealed a tight coupling between pupil diameter and neural responses in the noradrenergic locus coeruleus (Breton-Provencher & Sur, 2019; Joshi et al., 2016; Reimer et al., 2016; Liu et al., 2017; Varazzani et al., 2015), which is supported by recent human fMRI studies (de Gee et al., 2017; Murphy, O’Connell, O’Sullivan, Robertson, & Balsters, 2014). However, some of these studies also found unique contributions to pupil size in other subcortical regions, such as the cholinergic basal forebrain, dopaminergic midbrain, and the superior and inferior colliculi (de Gee et al., 2017; Joshi et al., 2016; Mridha et al., 2019; Reimer et al., 2016). Several lines of evidence reinforce a putative link between pupil diameter and the dopamine system, for example in patients with Parkinson’s disease (Kringelbach, Jenkinson, Owen, & Aziz, 2007; Manohar & Husain, 2015; Mathôt, 2018; Varazzani et al., 2015; Weinshenker & Schroeder, 2007), and although a link between dopamine and the P3 has also been observed in Parkinson’s patients, evidence is more mixed (see e.g. Bertram et al., 2020). The P3 has also been used as an electrophysiological correlate of feedback-evoked phasic catecholamine release in the cortex (Nieuwenhuis et al., 2005; Polich, 2007; Rangel-Gomez et al., 2013). Therefore, although these physiological markers of phasic arousal tend to co-occur, they may be driven by (partly) different sources. In line with this, we found that the behavioral impact of feedback-related pupil responses and the slow wave component of ERPs was also distinct: the slow wave ERP predicted behavioral speed-accuracy adaptation on the subsequent trial, which was absent for pupil size (Eckstein, Guerra-Carrillo, Miller Singley, & Bunge, 2017; Kamp & Donchin, 2015b; Murphy et al., 2011; LoTemplio et al., 2019). It is important to note, however, that different sources of noise contribute to pupil and EEG measurements, which might partly explain the absence of correlation between the two described here. For example, (i) participant fatigue, comfort, readiness, head movements, and eye movements all have a major impact on data quality but might differentially impact the EEG vs. pupil size signals, (ii) common additional sources of noise in EEG data include the cardiac signal (electrocardiogram, ECG) and movement artifacts caused by muscle contraction, e.g. in the neck (electromyogram, EMG), while luminance fluctuations, blinks and “pupillary hippus” contribute to noise in pupillometry data, and (iii) the respective acquisition systems introduce their own measurement noise. Understanding the relationship between changes in pupil dilation and the amplitude of the slow wave component and P3 component is an important avenue for future research.

Further, we show that feedback-related pupil responses and the slow wave component of ERPs only reflect surprise when feedback is fully visible. Although there is consensus that some perceptual and cognitive processes may unfold in the absence of awareness, it is highly debated which functions (if any) may need consciousness to emerge (Dehaene & Naccache, 2001; Hommel, 2007; Kunde et al., 2012; van Gaal et al., 2012). Many perceptual and cognitive processes may partly unfold unconsciously, typically demonstrated in masked priming studies, in which a task-irrelevant unconscious stimulus facilitates responding to a subsequent task-relevant conscious stimulus (Kiefer et al., 2011; Kiesel, Kunde, & Hoffmann, 2007; Lamme, 2010). These “simple” priming effects are typically explained by assuming that the fast feedforward sweep of neural processing is relatively unaffected by masking and is able to unconsciously affect ongoing behavioral responses (Dehaene & Changeux, 2011; van Gaal et al., 2012). We also observed here that the same masked stimuli used as feedback in the main experiment (the words error/correct) could induce behavioral priming when presented in the context of a masked priming task (**Fig. 3**). However, when the same stimuli were used as feedback stimuli in a perceptual decision task, no prediction error responses were observed (correctness x confidence interaction). Speculatively, error detection mechanisms (main effect of correctness) could still be observed when feedback was masked in the current task design (**Fig. 3**). Although evidence was statistically relatively weak, the main effects of correctness, especially in pupil size, were significant. This may not be overly surprising, because previous studies have shown that error detection mechanisms may unfold (at least partially) in the absence of error awareness (Charles et al., 2013; Cohen et al., 2009; Nieuwenhuis et al., 2001; Overbeek et al., 2005; Shalgi, 2012). However, our results revealed more importantly an absence of confidence x correctness interactions on low visibility feedback. This may suggest that to incorporate subjective confidence in feedback-driven prediction error computations, awareness of the decision outcome (feedback) is crucial. This may suggest that prediction error computation cannot rely on feedforward responses alone, in contrast to e.g. masked priming, and requires (bidirectional) interactions (i.e. recurrent processing) between higher-order and lower-order regions, a phenomenon mainly observed when stimuli are presented above the threshold of conscious perception (Dehaene & Changeux, 2011; van Gaal & Lamme, 2012). Further unraveling the underlying neural processes dissociating “objective error processing” from “prediction error” computation is important for further understanding the potential scope and limits of unconscious information processing.

Although we observed that confidence associated surprise was only present when the decision outcome (feedback) was presented fully consciously, previous studies have shown that pupil size is sensitive to *implicit* surprise and effort invested in a cognitive task. For example, it has recently been demonstrated that pupil size increases when the level of cognitive effort invested in a (conflict) task is high, even when subjects are not aware of systematic differences in difficulty between conditions (Diede & Bugg, 2017). Related, it has been shown recently that when agents are not aware of specific transitional rules in an implicit learning task, both the pupil and central ERP potentials (reminiscent of the mismatch negativity) may still signal surprise when statistical regularities in stimulus transitions are violated (Alamia et al., 2019; see also Meijs et al., 2018). Although intriguing, both tasks can be considered “implicit” (learning/conflict) tasks, because stimuli were always presented fully consciously and subjects were just not aware of differences in the probabilities of occurrence of specific stimuli. Therefore, these effects cannot be directly compared to situation in which stimulus visibility is reduced.

Although the main goal of this study was to test the association between putative measures of central arousal state (pupil response and the slow wave component of ERPs) and prediction error computation, we also explored the same association for other ERP components associated with feedback processing, such as the feedback related negativity (FRN) and the P3 (Cohen et al., 2011; Correa et al., 2018). Previously, using a probabilistic reversal learning task, we observed that the amplitude of the FRN was strongly linked to the signed prediction error variable (“objective prediction error”) derived from reinforcement learning modeling (Correa et al., 2018). This relationship was strongly attenuated when feedback awareness was reduced. Future work is needed to explore in more detail the relationship between different ERP components (e.g. FRN, P3, slow wave ERP) and specific aspects of prediction error computation and how these may be differentially affected by levels of (feedback) awareness.

## Acknowledgements

This study was supported by the Brazilian Science Without Borders program.

## Author contributions

JWdG, conceptualization, data curation, formal analysis, visualization, writing – original draft preparation, writing – review and editing; CC, conceptualization, investigation, writing – original draft preparation, writing – review and editing, funding acquisition; MW, formal analysis; TD, conceptualization, writing – review and editing; SvG, conceptualization, writing – original draft preparation, writing – review and editing, supervision, funding acquisition.

---

**Figure S1.**
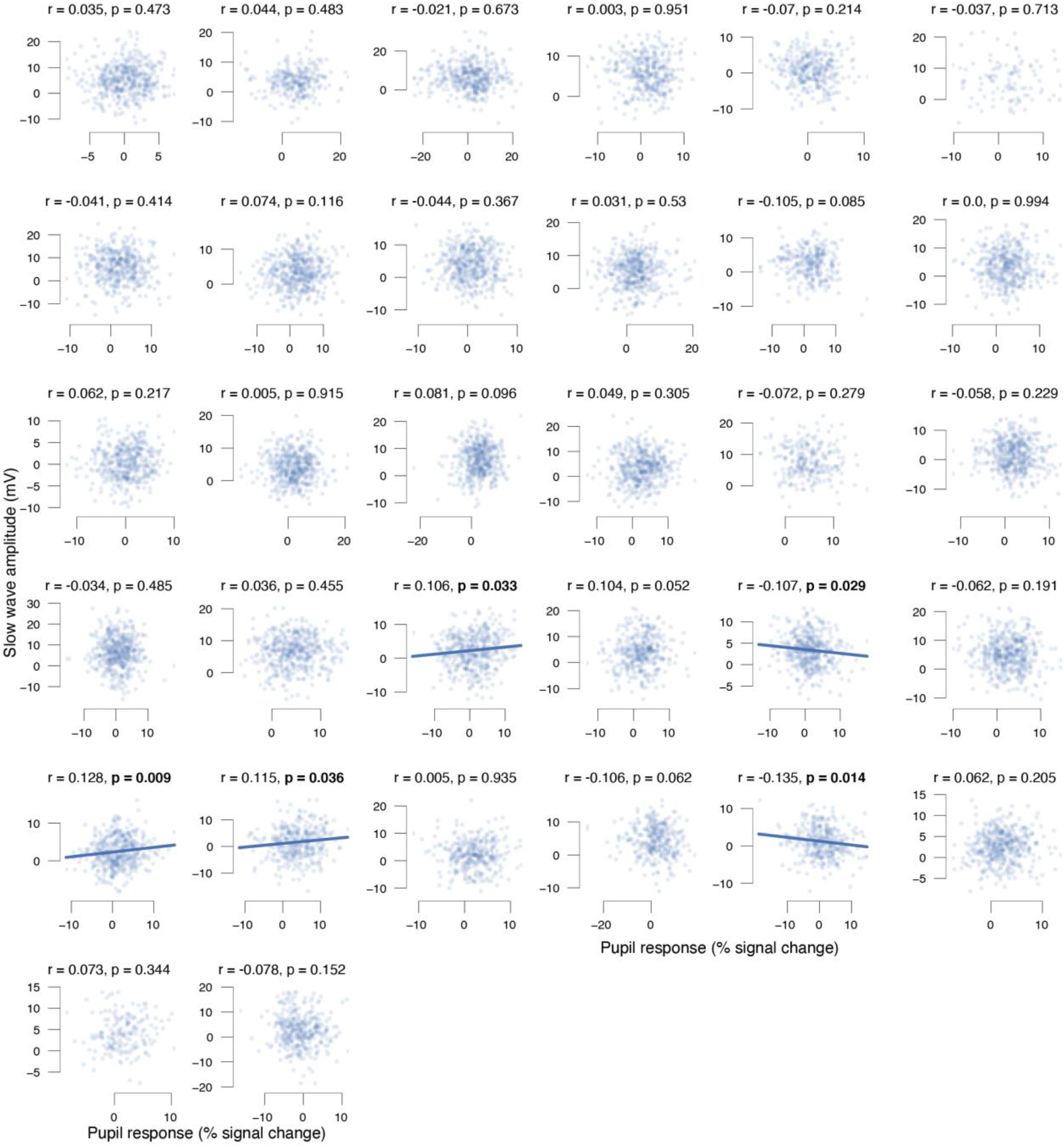
Trial-by-trial relationship between high visibility feedback-related pupil response and pre-trial baseline pupil size. Stats, Pearson correlation; data points, single trials; each panel is a different subject.

**Figure S2.**
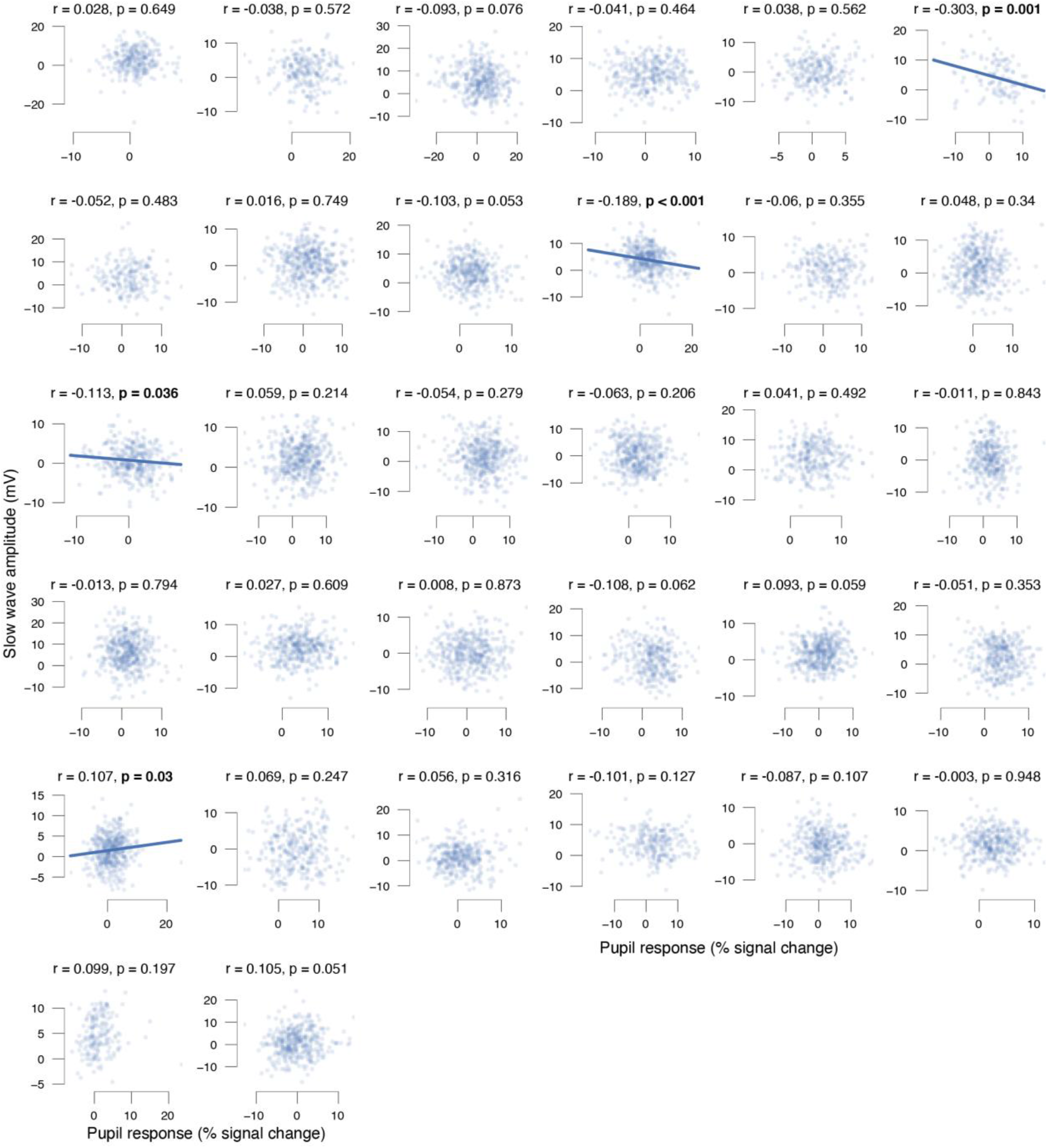
Trial-by-trial relationship between low visibility feedback-related pupil response and pre-trial baseline pupil size. Stats, Pearson correlation; data points, single trials; each panel is a different subject.

**Figure S3.**
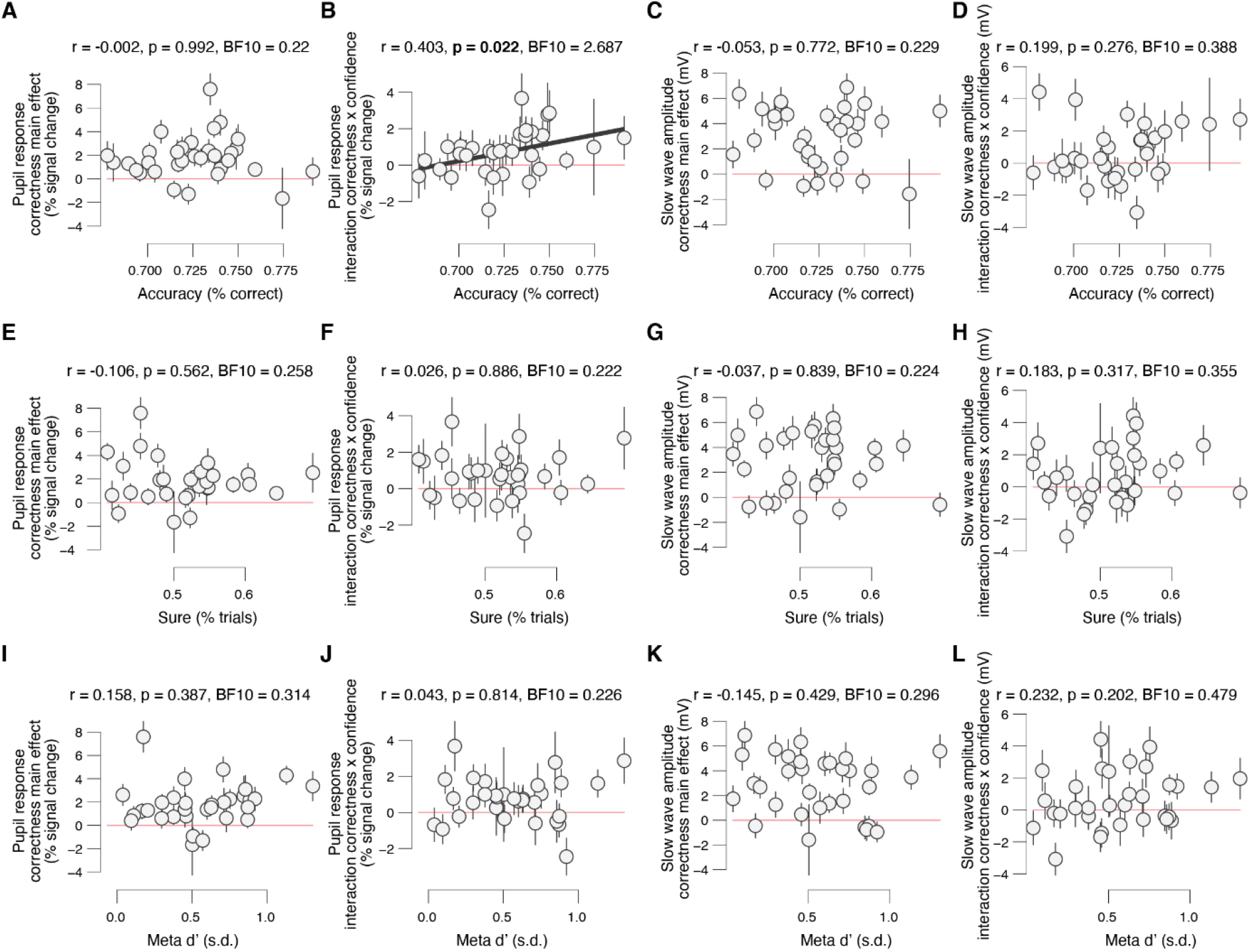
**(A)** High visibility feedback-related pupil response correctness main effect ([error/unsure + error/sure] - [correct/unsure + correct/sure]) plotted against overall accuracy. Stats, Pearson correlation; datapoints, individual subjects (N=32); error bars, 60% confidence intervals (bootstrap). **(B)** As A, but for correctness x confidence interaction effect ([error/sure - correct/sure] - [error/unsure - correct/unsure]). **(C,D)** As A,B, but for high visibility feedback-related slow wave amplitudes. **(E-F)** As A-D, but for overall confidence. **(I-L)** As A-D, but for overall meta-sensitivity.

**Figure S4.**
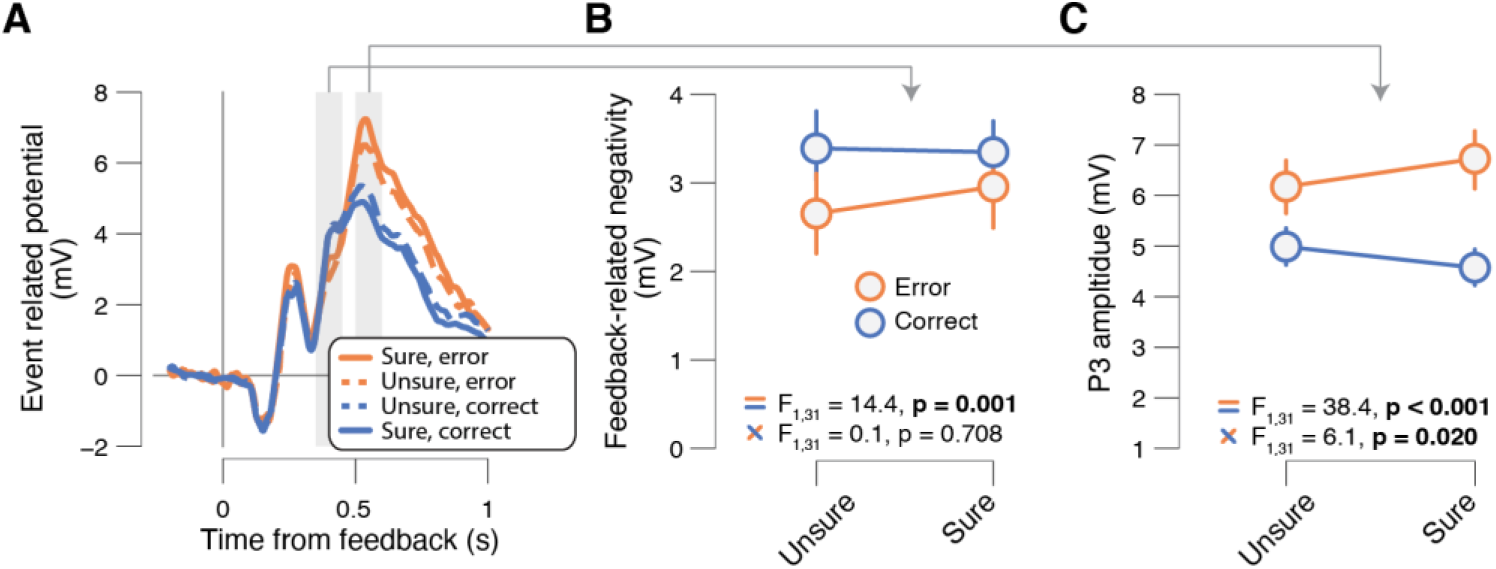
**(A)** High visibility feedback-related event-related potential (ERP) time courses, sorted by correctness (error, correct) and confidence (sure, unsure). Grey box, interval for averaging values on single trials. **(B)** High visibility feedback-related negativity (FRN) sorted by correctness (error, correct) and confidence (sure, unsure). Stats, ANOVA (Materials and Methods): top, correctness main effect; bottom, correctness x confidence interaction. **(C)** As B, but for P3 peak amplitudes.

**Figure S5.**
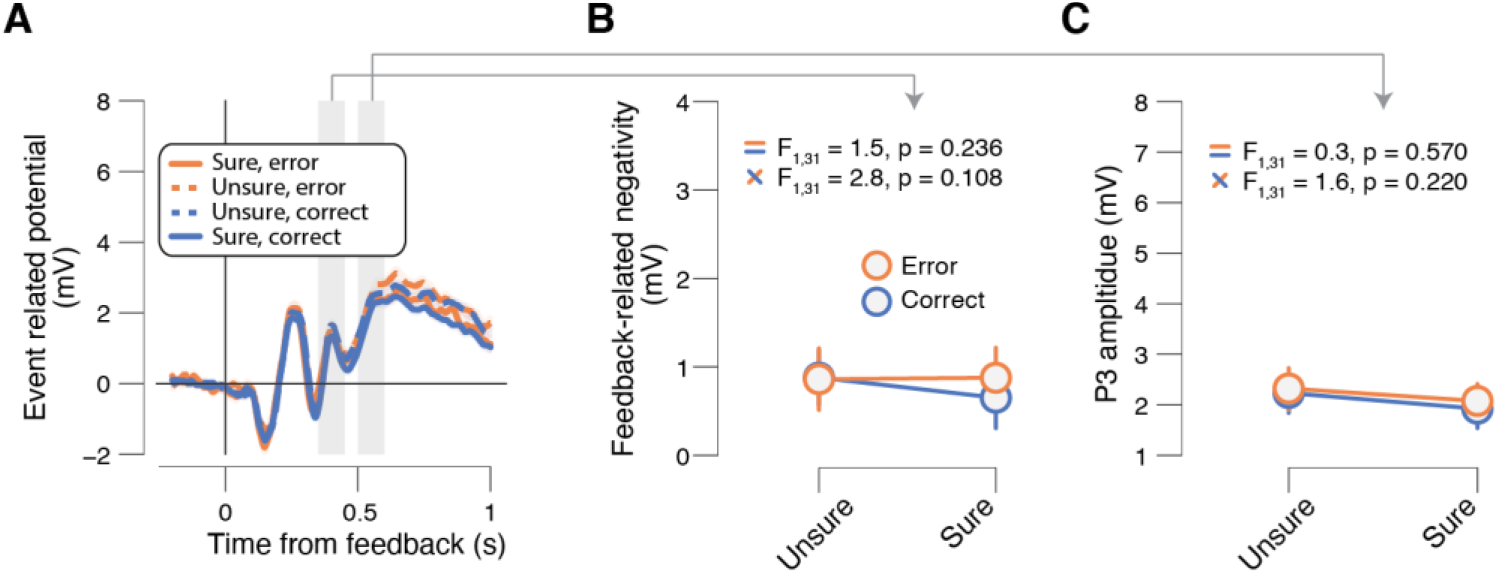
**(A)** Low visibility feedback-related event-related potential (ERP) time courses, sorted by correctness (error, correct) and confidence (sure, unsure). Grey box, interval for averaging values on single trials. **(B)** Low visibility feedback-related negativity (FRN) sorted by correctness (error, correct) and confidence (sure, unsure). Stats, ANOVA (Materials and Methods): top, correctness main effect; bottom, correctness x confidence interaction. **(C)** As B, but for P3 peak amplitudes.

**Figure S6.**
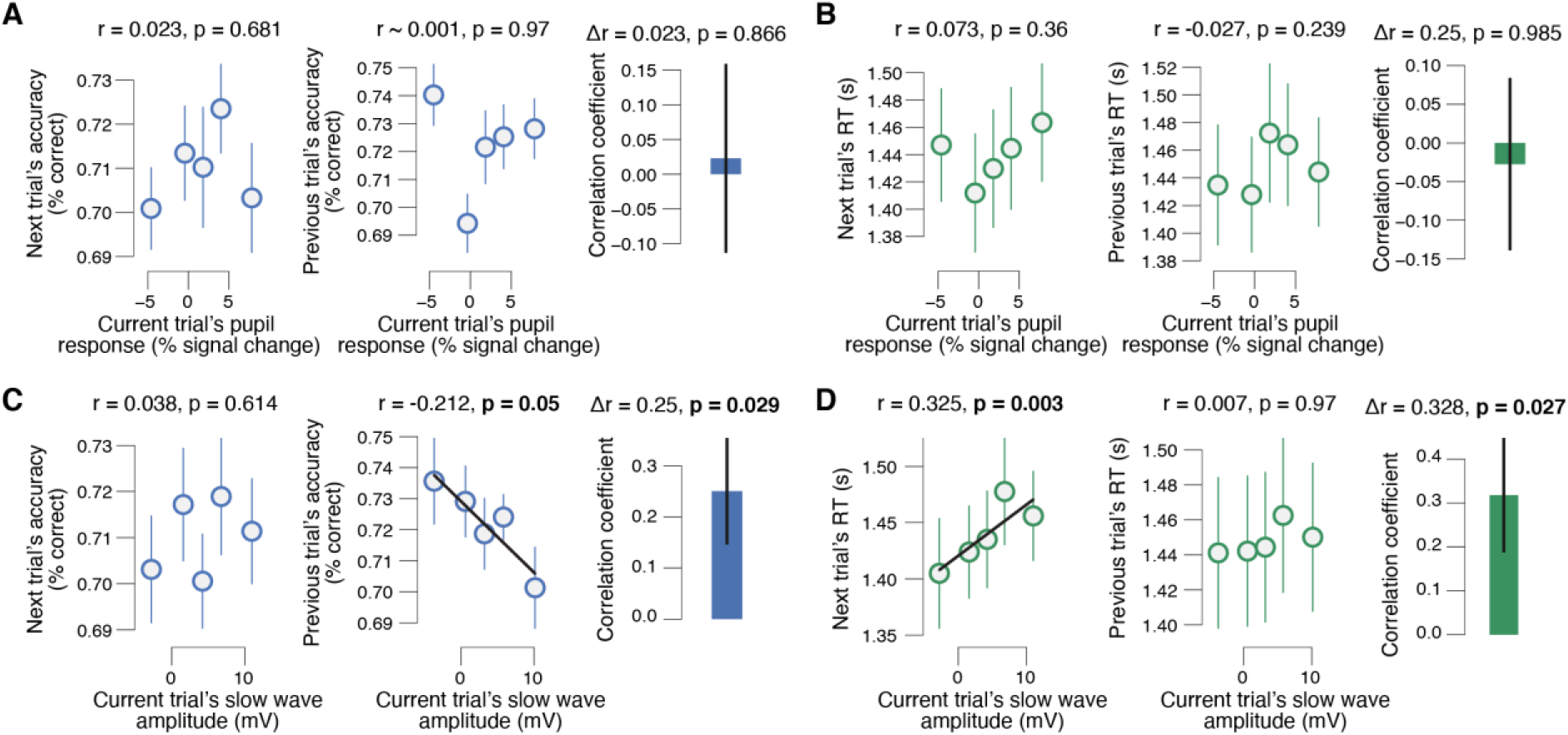
**(A)** Left: relationship between high visibility feedback-related pupil responses and accuracy on the next trial. Stats, Pearson correlation; error bars, s.e.m. across subjects; data is binned for visualization purposes only. Middle, as left, but for accuracy on the previous trial. Right: group average *difference* between correlation coefficients from left and middle panels. Error bar, s.e.m. across subjects. **(B)** As A, but for RT. **(C,D)** As A,B, but for high visibility feedback-relation slow wave amplitudes.

